# Feeding-structure morphogenesis in “rhabditid” and diplogastrid nematodes is not controlled by a conserved genetic module

**DOI:** 10.1101/2023.11.07.565949

**Authors:** Tobias Theska, Ralf J. Sommer

**Affiliations:** Max Planck Institute for Biology Tübingen (MPI-B), Department for Integrative Evolutionary Biology, Max-Planck-Ring 9, 72076 Tübingen, Germany

**Keywords:** morphological novelty, stoma development, evo-devo, *Caenorhabditis elegans*, *Pristionchus pacificus*

## Abstract

Disentangling the evolution of the molecular processes and genetic networks that facilitate the emergence of morphological novelties is one of the main objectives in evolutionary developmental biology. Here, we investigated the evolutionary history of a gene regulatory network controlling the development of novel tooth-like feeding-structures in diplogastrid nematodes. Focusing on NHR-1 and NHR-40, the two transcription factors that regulate the morphogenesis of these feeding structures in *Pristionchus pacificus*, we sought to determine whether they have a similar function in out-group nematode *Caenorhabditis elegans*, which has typical “rhabditid” flaps instead of teeth. Contrary to our initial expectations, we found that they do not have a similar function. While both receptors are co-expressed in the tissues that produce the feeding structures in the two nematodes, genetic inactivation of either receptor had no impact on feeding-structure morphogenesis in *C. elegans*. Transcriptomic experiments revealed that NHR-1 and NHR-40 have highly species-specific regulatory targets. These results suggest two possible evolutionary scenarios: either the genetic module responsible for feeding-structure morphogenesis in Diplogastridae already existed in the last common ancestor of *C. elegans* and *P. pacificus*, and subsequently disintegrated in the former as NHR-1 and NHR-40 acquired new targets, or it evolved in conjunction with teeth in Diplogastridae. These findings indicate that feeding-structure morphogenesis is regulated by different genetic programs in *P. pacificus* and *C. elegans*, hinting at developmental systems drift during the flap-to-tooth transformation. Further research in other “rhabditid” species is needed to fully reconstruct the developmental genetic changes which facilitated the evolution of novel feeding structures in Diplogastridae.

**Research Highlights:** Combining CRISPR-based mutagenesis, geometric morphometrics, and transcriptomics, we found that the genetic module governing the morphogenesis of novel feeding structures in diplogastrid nematodes is not conserved in the “rhabditid” *C. elegans*.

## Introduction

Evolutionary developmental biology (evo-devo) investigates how organismal development evolves and in which ways developmental processes guide organismal evolution, with the ultimate aim of mechanistically understanding phenotypic transformations and the emergence of new traits (Minelli, 2015). In evo-devo, morphological novelties are defined based on structural homology and relative to ancestral character states (Wagner, 2014). Currently, two different types of novelties are broadly recognized: discrete novelties and individualized novelties. Discrete novelties are new structures which have no homologous counterpart in an ancestral species; individualized novelties constitute major deviations from one or more pre-existing traits which progressively evolved a new qualitative dimension or functional capacity (Wagner, 2014; Müller, 2021).

Nematodes (roundworms) have historically been used as model organisms to study myriad facets of biology and evolution (Corsi *et al*., 2015; Sommer, 2015), given the invaluable characteristics of some species for functional genetics. Indeed, selected nematode model organisms mainly reproduce as self-fertilizing hermaphrodites and therefore produce isogenic populations. In addition, many nematodes have a stringent mode of development in which a small and invariable number of cells generate the body, a phenomenon known as eutely. These features made it possible to homologize the worm’s anatomical traits down to the level of individual cells (Haag *et al*., 2018) rendering eutelic model nematodes, such as the “rhabditid” *Caenorhabditis elegans* and the diplogastrid *Pristionchus pacificus,* promising subjects for comparative evo-devo studies of morphological character transformations.

Despite some superficial similarities in their simplified cylindrical body shape, nematodes are far from morphologically uniform. A closer look reveals a stunning diversity in cuticular feeding structures (Wright, 1976; Malakhov, 1994; Susoy *et al*., 2015), which arguably explains how these microscopic animals were able to conquer nearly all ecosystems. When referring to the roundworm’s mouth (stoma), nematologists traditionally distinguish between the buccal cavity (the lumen of the mouth) and the buccal capsule (the inner cuticular lining of the mouth) (Fürst von Lieven & Sudhaus, 2000). Based on the underlying cells which secrete it, the buccal capsule can be subdivided into individual cuticular feeding structures (from anterior to posterior): cheilostom, gymnostom, prostegostom, mesostegostom, metastegostom, and telostegostom (De Ley *et al*., 1995). The cheilostom is secreted by skin cells (hypodermis) and is continuous with the outer body-wall cuticle of the worm. The gymnostom, which forms the middle part of the buccal capsule, was found to be associated with the anterior and posterior arcade syncytia (De Ley *et al*., 1995; Burr & Baldwin, 2016) - nematode-specific cells with enigmatic functions. The four posterior-most feeding structures are collectively referred to as the stegostom - the part of the buccal capsule that is made by pharyngeal cells. The prostegostom and mesostegostom are each produced by a set of “e” cells (pharyngeal epithelia); the metastegostom and telostegostom are each secreted by a set of “pm” cells (pharyngeal muscles) (De Ley *et al*., 1995; Burr & Baldwin, 2016; Harry *et al.,* 2022). Interestingly, all of these elements are cuticular; they, just like the body-wall cuticle, constantly have to be remodeled, degraded and re-secreted, during the worm’s periodic larval molts.

Some of these feeding structures differ drastically between the closely related “rhabditid” and diplogastrid nematodes (Figure 1a,b,d). “Rhabditids” represent a paraphyletic group (Figure 1e) whose members typically have elongated buccal capsules in which cheilo-, gymno-, pro-, and mesostegostoms form a rigid cuticular tube (Figure 1a). Their metastegostoms are shaped into simple cuticular triangles called flaps (Figure 1a,e) (Fürst von Lieven & Sudhaus, 2000). These structures are not actively moveable and likely function as valves, which prevent regurgitation of bacterial foods from the pharynx to the buccal cavity. In contrast, in Diplogastridae, a monophyletic clade that emerged from the paraphyletic “rhabditids” (Figure 1e), buccal capsules seldomly come in the form of rigid tubes and flaps do not exist (Fürst von Lieven & Sudhaus, 2000; Sudhaus, 2014). Instead, cheilostom and gymnostom are developed as cuticular rings which are telescoped into each other, giving the buccal capsule a barrel-like shape (Figure 1b,d). Strikingly, the metastegostom of diplogastrids evolved into large hook-shaped teeth (Figure 1b,d,e) that can actively be moved via pharyngeal muscle contractions and used for predatory feeding (Figure 1c) (Fürst von Lieven & Sudhaus, 2000; Susoy *et al*., 2015; Sommer, 2015). Their pro- and mesostegostom evolved into slender and flexible hinges, which facilitate these movements relative to the remainder of the buccal capsule (compare Figure 1a and 1b) (Harry *et al*., 2022).

**Figure 1.**
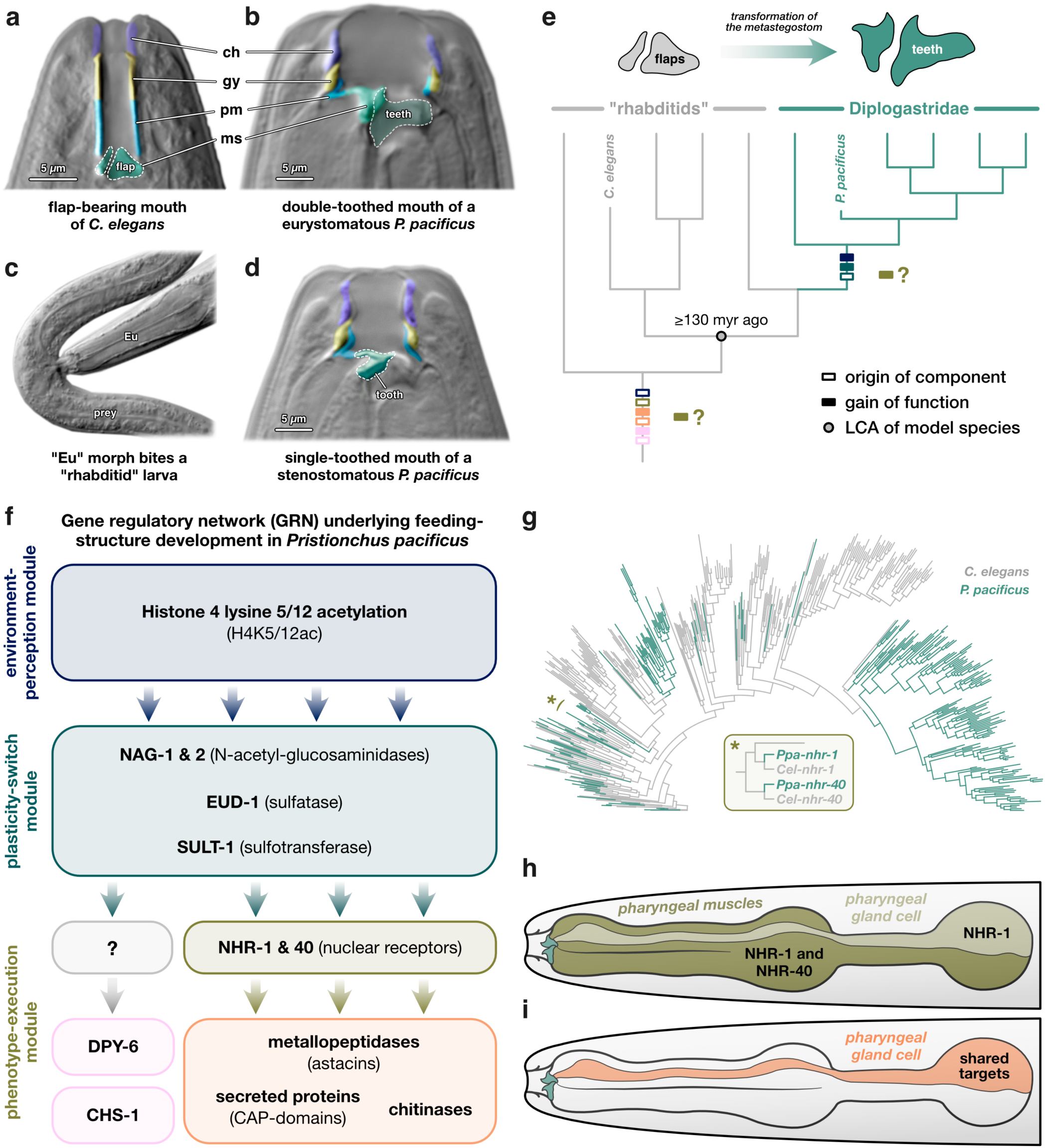
Evolutionary history and developmental genetics of feeding structures in “rhabditids” and Diplogastridae. Microscopic images (with DIC) of the mouths of (a) *C. elegans*, (b) eurystomatous *P. pacificus*, (d) stenostomatous *P. pacificus*. Homologous feeding structures are correspondingly color-coded (ch = cheilostom, gy = gymnostom, pm = promesostegostom, ms = metastegostom). (c) Microscopic images (with DIC) of a eurystomatous *P. pacificus* preying on a “rhabditid” larva. (e) Phylogeny depicting the flap-to-tooth transformation of the metastegostom during the “rhabditid”-to-diplogastrid transition. Time estimate for LCA is based on Howard *et al*. (2022). Color coding of the boxes on the branches of the phylogeny indicates the origins of components of (f) the modular GRN controlling feeding-structure development in *P. pacificus*. Empty boxes in (e) correspond to the origin of the molecular players; filled boxes indicate the gain of their functions related to feeding-structure development (LCA = last common ancestor). (g) Phylogeny of all *nhr* genes encoded in the genomes of *C. elegans* and *P. pacificus* recreated from the data in Sieriebriennikov *et al*. (2020). The asterisk (*) indicates the location of the branch that harbors the one-to-one orthologs of NHR-1 and NHR-40. (h, i) Schematic representation of the expression patterns of NHR-1 and NHR-40 and their shared regulatory target genes in *P. pacificus*. Depicted is the head region in lateral position; the buccal cavity is on the left.

Given nematode eutely, the individual cells which secrete the buccal capsule can be homologized. Thus, one can also directly homologize the individual feeding structures they produce, including the metastegostomatal teeth of diplogastrids and the flaps of the paraphyletic “rhabditids” (Figure 1a,b,e). A recent study reconstructed the buccal cavity of *P. pacificus* and showed that their feeding structures are indeed homologous to the ones in *C. elegans* (Harry *et al*., 2022). Nonetheless, the movable teeth of *P. pacificus* and other diplogastrids are unlike any other feeding structure in nematodes. While they evolved from a pre-existing structure (the metastegostom), they have reached an extreme point of phenotypic deviation from its ancestral state (flaps); and thereby developed into uniquely specialized traits that gained the quality of raptorial structures (hook-shaped teeth), which now serve a function their precursors did not, predation. Hence, the diplogastrid’s teeth are an example of individualized novelty. Together, the fact that the evolutionary history of this trait can be reconstructed unambiguously (as robust phylogenetic backbones are available), and that these nematodes readily lend themselves for functional genetic experiments, renders the flap-to-tooth transformation that accompanied the “rhabditid”-to-diplogastrid transition a model system for studies on the molecular and developmental underpinnings of morphological novelties.

Revealing the molecular components of the gene regulatory network (GRN) which controls the development of diplogastrid feeding structures was a major focus of many recent genetic studies (for reviews see, Sommer *et al*., 2017; Sommer, 2020). This is particularly due to the fact that most diplogastrids display a feeding polyphenism: in response to various environmental inputs such as pheromone signals (Werner *et al*., 2018) or compositional differences in the media surrounding them (Werner *et al*., 2017), these worms can develop either into narrow-mouthed bacterivores (stenostomatous morph; “St”) or wide-mouthed predators (eurystomatous morph; “Eu”) (Susoy *et al.,* 2015). In *P. pacificus*, both morphs carry a moveable metastegostomatal tooth at the anterior tip of their dorsal pharynx (Figure 1b,d). However, Eu morphs possess a second cuticular tooth at the anterior tip of the right subventral pharynx (Figure 1b), where St animals only possess a flat cuticle ridge (Fürst von Lieven & Sudhaus, 2000; Sommer, 2015; Theska *et al*., 2020, Harry *et al*. 2022). By moving these two teeth against each other, Eu worms can rupture the cuticle of their prey (Figure 1c).

Work over the last decade has established that a modular GRN governs the development of these phenotypically plastic feeding structures (Figure 1f). In the center of this network sits a plasticity-switch module that controls which of the two alternative feeding morphologies will be developed (Sommer, 2020). It consists of an X-chromosomal multi-gene locus that is expressed in the central nervous system and the pharynx of the worm (Sieriebriennikov *et al*., 2018). Upstream of the plasticity-switch module is an environment-perception module, which governs the activity of the plasticity-switch module in correspondence with environmental signals perceived during early larval development. This module works largely through the epigenetic mechanism of histone 4 lysine K 5/12 acetylation (H4K5/12ac) (Werner *et al.,* 2023). The GRN is completed by a phenotype-execution module that is downstream of the plasticity switch and controls the morphogenesis of the feeding structures (Sommer, 2020). The core of this module are two transcription factors (TFs), the nuclear receptors NHR-1 and NHR-40, which control the expression of multiple protein classes involved in the modification and degradation of extracellular matrix and cuticle - the materials that make up the worm’s feeding structures (Kieninger *et* al., 2016; Sieriebriennikov *et al*., 2020) (Figure 1f). Recent studies also identified the mucin-type protein DPY-6 and the chitin synthase CHS-1 as members of this module (Sun & Theska *et al*., 2022; Sun *et al*., 2023). DPY-6 is the first known protein constituent of the buccal capsule and located mostly in the cheilostom. CHS-1 synthesizes chitin, and *Ppa-chs-1* mutants are teethless, showing for the first time that chitin is a structural constituent of nematode feeding structures (Sun *et al*., 2023). However, the transcription factors, which regulate the expression of these cuticle-synthesis related components of the phenotype-execution module, have not yet been identified (Figure 1f).

Interestingly, the evolutionary histories of the genes establishing the *P. pacificus* mouth-form GRN are diverse. All known components of the plasticity-switch module are derived from recent gene-duplication events, which accompanied the flap-to-tooth transformation during the “rhabditid”-to-diplogastrid transition (Figure 1e,f) (Sieriebriennikov *et al*., 2018; Sommer, 2020). As such, these young genes seemed to have gained their functions related to feeding-structure development immediately or early in their evolutionary history (Figure 1e). This is in stark contrast to some of the components of the phenotype-execution module. Both DPY-6 and CHS-1 are deeply-conserved proteins with functions related to buccal-capsule development, which predate the “rhabditid”-to-diplogastrid transition (Figure 1e) (Sun & Theska *et al*., 2022; Sun *et al*., 2023). Additionally, both NHR-1 and NHR-40 exist as one-to-one orthologs in *C. elegans* and *P. pacificus* (Figure 1g) and therefore already existed in their last common ancestor (LCA) (Figure 1e) (Sieriebriennikov *et al*., 2020). Yet, the target genes that these receptors co-regulate to mediate feeding-structure development in diplogastrids are fast evolving (Sieriebriennikov *et al*., 2020), even though the classes of cuticle-modifying proteins to which they belong (astacins, chitinases, and cysteine-rich secreted proteins) are ancient (Figure 1e). This contrast is even more stunning given that, in nematodes, NHRs are rapidly evolving, too (Robinson-Rechavi *et al*., 2005; Sural & Hobert, 2021). Most of the *nhr* genes in the genomes of *C. elegans* and *P. pacificus* are entirely lineage-specific and only a small handful of them (like *nhr-1* and *nhr-40*) is conserved between these worms (Figure 1g) (Sieriebriennikov *et al*., 2020). In *P. pacificus*, both of these receptors are expressed in the pharyngeal muscles, including the specific cells that secrete the metastegostomatal teeth (Figure 1h). Additionally, *nhr-1* is expressed in the dorsal pharyngeal gland (Sieriebriennikov *et al*., 2020), which, in diplogastrids, is a massively enlarged gland cell that runs through the pharyngeal muscles, connects to the metastegostom, and opens into the buccal cavity via a hole in the dorsal tooth (Figure 1h) (Riebesell & Sommer, 2017). Interestingly, all shared regulatory targets of NHR-1 and NHR-40 in *P. pacificus* are co-expressed in that precise cell, too (Figure 1i) (Sieriebriennikov *et al*., 2020).

In light of the fact that the teeth of diplogastrids are individualized novelties that evolved from “rhabditid” flaps, we were eager to ask: does the entire phenotype-execution module - including the NHRs regulating cuticle-modifying enzymes - represent an ancient genetic cassette that is also functionally conserved as a mouth-development program in “rhabditids”, such as *C. elegans*? This would imply that the feeding-structure GRN of diplogastrids evolved by building a new plasticity-switch module (which is made of evolutionarily young genes) upon an old mouth-development program. Alternatively, the two old NHRs may control a different biological function in “rhabditids”. In this case, they might have been co-opted into a new regulatory context - that is borrowed for stoma morphogenesis - in diplogastrids. To address these possibilities, we set out to study whether these nuclear receptors are expressed in the stomatal tissues of “rhabditid” model nematode *C. elegans* and whether their functional loss would cause developmental aberrations in the mouth of these worms.

## Material & methods

### Nematode husbandry

Standard protocols for the maintenance of laboratory cultures of rhabditid nematodes (Stiernagle, 2006) were followed. Worms were cultured on 6cm Petri dishes with nematode growth medium (NGM). 300µl of *Escherichia coli* (OP50) were provided as a food source. Culture plates were stored at 20°C.

### Light microscopy with differential interference contrast (DIC)

For microscopy, all specimens were mounted on object slides with 5% Noble Agar pads containing sodium azide (0.5% NaN_3_) and subsequently examined using a Zeiss Axio Imager.Z1 microscope with a Zeiss Plan-Apochromate 100 x 1.4 DIC objective. Image stacks (Grayscale, 16-bit) were taken using a monochromatic Zeiss Axiocam 506 CCD camera. The Zen 2 Pro Software (version 2.0.14283.302) was used for digital microscopy and image acquisition.

### CRISPR/CAS9 mutagenesis

We used the CRISPR/CAS9 system to create mutations in the coding sequences of *Cel-nhr-1* and *Cel-nhr-40*, specifically targeting their DNA-binding domains. We followed standard mutagenesis protocols for nematodes (Ghanta *et al*., 2021). Target-specific CRISPR RNAs (crRNAs) were generated based on the genome sequence of *C. elegans* (www.wormbase.org). The CAS9 protein, the universal trans-acting CRISPR RNA (tracrRNA), and the crRNAs were ordered from IDT (Alt-R product line). Ribonucleoprotein complexes (RNPs) were created by combining 0.5µl of CAS9 nuclease (10µg/ml stock) with 5µl of tracrRNA (0.4µg/µl stock) and 2.8µl of crRNA (0.4µg/µl stock). This mix was incubated at 37°C for 15min and subsequently cooled to room temperature. The RNP mix was diluted by adding enough Tris-EDTA buffer to bring the total volume of the mix to 20µl. In order to avoid clogging in the injection needles, the final mix was centrifuged at 14,000rpm for 2min. Injections were conducted with an Eppendorf FemtoJet microinjector and an inverse Zeiss AxioVert microscope coupled to an Eppendorf TransferMan micromanipulator. RNP complexes were injected into the distal gonadal rachis of young adult hermaphrodites. Injected P0 animals were isolated and allowed to lay eggs for 24h at 20°C. After that, P0s were removed from the plates and F1 animals were left to develop. Upon reaching maturity, F1s were singled to fresh plates, allowed to lay eggs for 24h, and subsequently genotyped. DNA of candidate F1 animals was recovered via single worm lysis (SWL), target sites were PCR amplified (using the Taq PCR Master Mix produced by Qiagen), and candidate mutations were identified using Sanger sequencing (performed by Azenta Life Sciences). If heterozygous F1 mutants were found, their F2 offspring were genotyped in a similar fashion to identify homozygous mutant carriers. Multiple homozygous mutant lines were isolated, including frameshift mutants with premature stop codons. Backups of these homozygous mutant strains were stored at -80°C and can be provided by the Sommer lab upon request.

### Transgenesis and SEC-based protein tags

Our CRISPR-based transgenesis experiments largely followed the well-established SEC approach (Dickinson *et al.,* 2015) with minor modifications. In short, we used the original pDD268 plasmid to create a transcriptional reporter for *Cel-nhr-1* (pre-heatshock) and an N-terminally mNeonGreen(mNG)-tagged version of *Cel-*NHR-1 (post-heatshock). pDD268 was digested overnight with ClaI and SpeI in a thermocylcer set to 37°C. We excluded the native start codon of *Cel-nhr-1* from the 3’-homology arm in the donor plasmid in order to reduce the likelihood of internal splicing initiation in the mutant strain that expresses the mNG-tagged version of *Cel*-NHR-1 (RS3881). We used the InVitrogen HiFi Miniprep Kit (including the optional wash step with Qiagen PB buffer) to extract plasmids from 4ml bacterial overnight cultures. Digested SEC-plasmids and PCR-amplified homology arms were cleaned with the Qiaquick MinElute kit, assembled into donor plasmids with NEB’s DNA HiFi Assembly Kit, and subsequently transformed into *E. coli* (NEB 5-alpha). We injected donor plasmids (20 ng/µl) together with prepared RNP complexes. Fluorescent worms were imaged using a Leica SP8 confocal laser-scanning microscope (cLSM).

### Quantification of mouth-form differences among wild-type and mutant worms

We quantified morphological differences in the mouths of wild-type and mutant worms, using our recently published protocol (Theska et al., 2020) which combines two-dimensional landmark-based geometric morphometrics (GMM) with model-based clustering. We obtained image stacks of the nematode mouths in lateral position and recorded the X and Y coordinates of 15 fixed landmarks which capture all feeding structures using FIJI (ver. 2.1.0) (Schindelin et al., 2012). All steps of the GMM analysis were performed in R (ver. 4.3.0) (R Core Team, 2023) using a combination of the GEOMORPH (ver. 4.0.5) (Adams *et al*., 2013; Baken *et al*., 2021), MORPHO (ver. 2.11) (Schlager, 2017), and MCLUST (ver. 6.0.0.) packages (Scrucca *et al*., 2016). General Procrustes Analysis (GPA) was performed with the *gpagen* function of GEOMORPH, by minimizing Procrustes distances. We generated landmark data for 150 animals (50 per strain; three strains: N2, RS3635, RS3682), performed GPA, and subsequently checked for Procrustes distance outliers, using the *plotOutliers* function of GEOMORPH. Seven outliers were removed from the data for all subsequent steps of analysis. The outlier-corrected data set (n=143) contained landmark configurations from 44 wild-type animals (N2), 49 *Cel-nhr-1(tu1424)* mutant animals (RS3635), and 50 *Cel-nhr-40(tu1464)* mutant animals (RS3682). GPA was performed on the outlier-corrected data to obtain a shape matrix, to which we appended the log-transformed centroid size in order to generate a form data set (*sensu* Mitteroecker *et al.,* 2004). We subsequently performed a principal component analysis (PCA) on the form data set to visualize potential morphological differences among wild-type and mutant worms in a form space. Lastly, we assessed whether mutant worms could be classified as such and distinguished from wild-type worms, based on their mouth morphology. For this, we performed model-based clustering with the *Mclust* function of MCLUST. We only used “meaningful” principal components (mPCs) of form variation (identified with MORPHO’s *getMeaningfulPCs* function) as input variables for clustering, in order to avoid overparameterization (Theska *et al*., 2020). Clustering results were visualized by coloring each specimen according to cluster membership.

### RNA-seq experiments

To facilitate RNA-seq experiments on our *Cel-nhr-1(tu1424)* and *Cel-nhr-40(tu1464)* mutants being directly comparable to the RNA-seq experiments our lab previously conducted on *Ppa-nhr-1* and *Ppa-nhr-40* mutants, we adopted the sample collection approach of Sieriebriennikov *et al*. (2020) with minor modifications. Animals were collected for RNA extraction at 33h (mostly L2 and a few L3 larvae), 44h (L3 and L4 larvae), and 58h (young adults) after bleaching; the presence and expected distribution of the desired stages at each time point was verified prior to sample collection by screening 50 specimens on each plate under a dissecting microscope. Total RNA was extracted using the Direct-Zol RNA Mini prep kit (Zymo Research) according to the instructions provided by the manufacturer. We combined 500ng of RNA extracted at the 33h time point with 500ng of RNA extracted at the 44h time point. This sample type thus contains a total of 1µg RNA extracted from L2-4 larvae. We gathered 1µg of RNA extracted at the 58h time point (adult worms) and did not mix it with RNA extracted from any other time point. Therefore, we ended up with two sample types (”larval” and “adult”), which we shipped to Novogene for library preparation and mRNA sequencing. Libraries were sequenced as 150bp paired end reads on Illumina’s NovaSeq6000 platform. We sequenced the mRNA of three strains: the wild-type *C. elegans* strain N2, the *Cel*-*nhr-1* frameshift mutant RS3635(*tu1424*), and the *Cel*-*nhr-40* frameshift mutant RS3682(*tu1464*). Two biological replicates were collected for each of the larval and adult samples of each of these three strains. The obtained raw reads have been deposited in the European Nucleotide Archive (ENA) under the study accession number PRJEB68220.

### Quantification of transcripts and differential gene expression (DGE) analysis

Transcript abundances were estimated from raw read files relative to the *C. elegans* transcriptome using SALMON (ver. 1.5.2) (Patro *et al*., 2017; Srivastava *et al*., 2020) [84,85]. Using the entire *C. elegans* reference genome as a decoy, we built a decoy-aware transcriptome and indexed it for subsequent read mapping using an auxiliary *k*-mer hash (k=31). Reads were quantified against this index using Salmon’s *quant* command by running the program in mapping-based mode with selective alignment as mapping strategy. We allowed Salmon to infer the library type, and to learn and correct for fragment-level GC-biases and sequence-specific biases. Differential expression analyses (DEA) were carried out in R using BIOCONDUCTOR (ver. 3.17) (Gentleman *et al*., 2004), TXIMPORT (ver. 1.28.0) (Love *et al*., 2015), and DESEQ2 (ver. 1.40.1) (Love *et al*., 2014). Transcript-level abundance estimates generated by SALMON were summarized into gene-level count matrices using the *tximport* function. A DESeq data set was created with the *DESeqDataSetFromTximport* function and pre-filtered for transcripts which had at least ten counts across all samples. Stage-specific mutation effects on transcript abundances were modeled with the *DESeq* function. Obtained maximum likelihood estimates (MLE) of the log2-fold changes (L2FC) were shrunken with the *lfcShrink* function using the adaptive shrinkage estimator from the ASHR package (Stephens, 2017). Resulting minimum mean squared error (MMSE) estimates of the L2FC were reported for this study and used to visualize candidate target genes. We always included a L2FC-threshold of > 0.585 (absolute value) and an *s*-value threshold of < 0.001 into our hypothesis tests for differential expression. This is equivalent to asking whether there is sufficient evidence in the data that a given gene shows at least a 50% increase or decrease in expression levels, and that the direction of the observed effect was correctly estimated. Both as proof-of-concept and to facilitate comparability of datasets, we also used this pipeline to reanalyze the transcriptome data of wild-type *P. pacificus* (PS312), as well as *Ppa-nhr-1(-)* and *Ppa-nhr-40(-)* mutants obtained by Sieriebriennikov *et al*. (2020). Results of the DEAs can be found in Sup. Data 2 and 3.

### Pfam domain prediction, WormCat annotation, and overrepresentation analyses (ORA)

Protein domains encoded in the proteome of *C. elegans* were predicted using HMMER (ver. 3.3.2) (Mistry *et al.,* 2013) in conjunction with the Pfam-A.hmm database (ver. 3.1b2) (Finn *et al.,* 2016) (hmmscan, e-value cutoff: < 0.001). WormCat (ver. 2) annotations were used to explore biological functions of target genes (Holdorf *et al*., 2020). ORTHOFINDER (default mode; ver. 2.5.4) (Emms & Kelly, 2019) was used to define orthogroups between *P. pacificus* and *C. elegans* proteins, based on which WormCat annotations for *P. pacificus* were created. ORAs were based on Fisher’s Exact test; WormCat terms were considered overrepresented if FDR-corrected *P*-values were below a type I error rate of 5% (*P* < 0.05). Results of the ORAs can be found in Sup. Data 4 and 5.

### Data processing, data visualization, and illustration

All data was processed, formatted, and wrangled in R (ver. 4.3.0) (R Core Team, 2023) using the TIDYVERSE (Wickham *et al*., 2019). All scientific plots were created using GGPLOT2 (ver. 3.4.2) (Wickham *et al*., 2019). In order to make our plots accessible to people with color-vision deficiencies or color blindness, we used the scientifically-derived colormaps provided by the SCICO (ver. 1.4.0) and VIRIDIS (ver. 0.6.3) packages (Pedersen & Crameri, 2023; Garnier *et al*., 2023). Microscopic images were edited and adjusted for color, levels, lightness, and contrast in Affinity Photo (ver. 1.10.5). Scientific illustrations and final figures were created in Affinity Designer (ver. 1.10.5).

## Results

### *Cel-nhr-1* is broadly expressed in somatic tissues including pharyngeal muscles

First, we wondered whether *C. elegans’s* copies of *nhr*-1 and *nhr*-40 are expressed in the same tissues in which they are expressed in *P. pacificus*. Previous studies have shown that *Cel-nhr-40* is indeed expressed in the anterior pharyngeal muscles of *C. elegans*, as well as other tissues including head neurons, skin, and body wall muscles (Brožová *et al*., 2006; Taylor *et al*., 2021). Thus, the currently available data suggests that the expression of *nhr-40* in the pharynx - the tissue which secrets the metastegostom - is evolutionarily conserved across “rhabditids” and diplogastrids. We speculated that the same holds true for *nhr-1*, which would indicate that both receptors are co-expressed in the mouth-associated tissues in flap-bearing “rhabditid” nematodes. To verify this, we studied the expression of *nhr-1* in *C. elegans* via the creation of transcriptional and translational reporter lines.

Our transcriptional reporter lines revealed that *Cel-nhr-1* is broadly expressed in multiple somatic tissues at varying intensities. We found high expression levels in the entire skin (hypodermis) and body wall musculature from head to tail, as well as in the majority of the worm’s pharyngeal muscles (Figure 2a). The latter includes pm3-5, the cells which make up most of the muscular portions of the procorpus, metacorpus, and isthmus (Figure 2b). Relatively lower expression levels were found in the stomatal tissues (aa, ap, e1, e3, pm1, pm2), the terminal bulb of the pharynx (pm6-8), the intestine, and some ventral aspects of the somatic gonad (Figure 2a,b,c). Furthermore, animals carrying the translational reporter express a fluorescently labeled version of NHR-1 that is restricted to the nuclei of the cells in which *Cel-nhr-1* is expressed (Figure 2d). Specifically, using these localization patterns, we were able to identify the nuclei of the stomatal and pharyngeal muscle cells for which we already have seen expression signals in the transcriptional reporter line (Figure 2b,c). These results confirm the presence of *Cel-*NHR-1 protein in exactly those cells that secrete the posterior buccal capsule (Figure 2d). Additionally, we found *Cel-*NHR-1 in the nuclei of multiple nerve-ring neurons, possibly including amphid-associated neurons or glia cells (Figure 2d).

**Figure 2.**
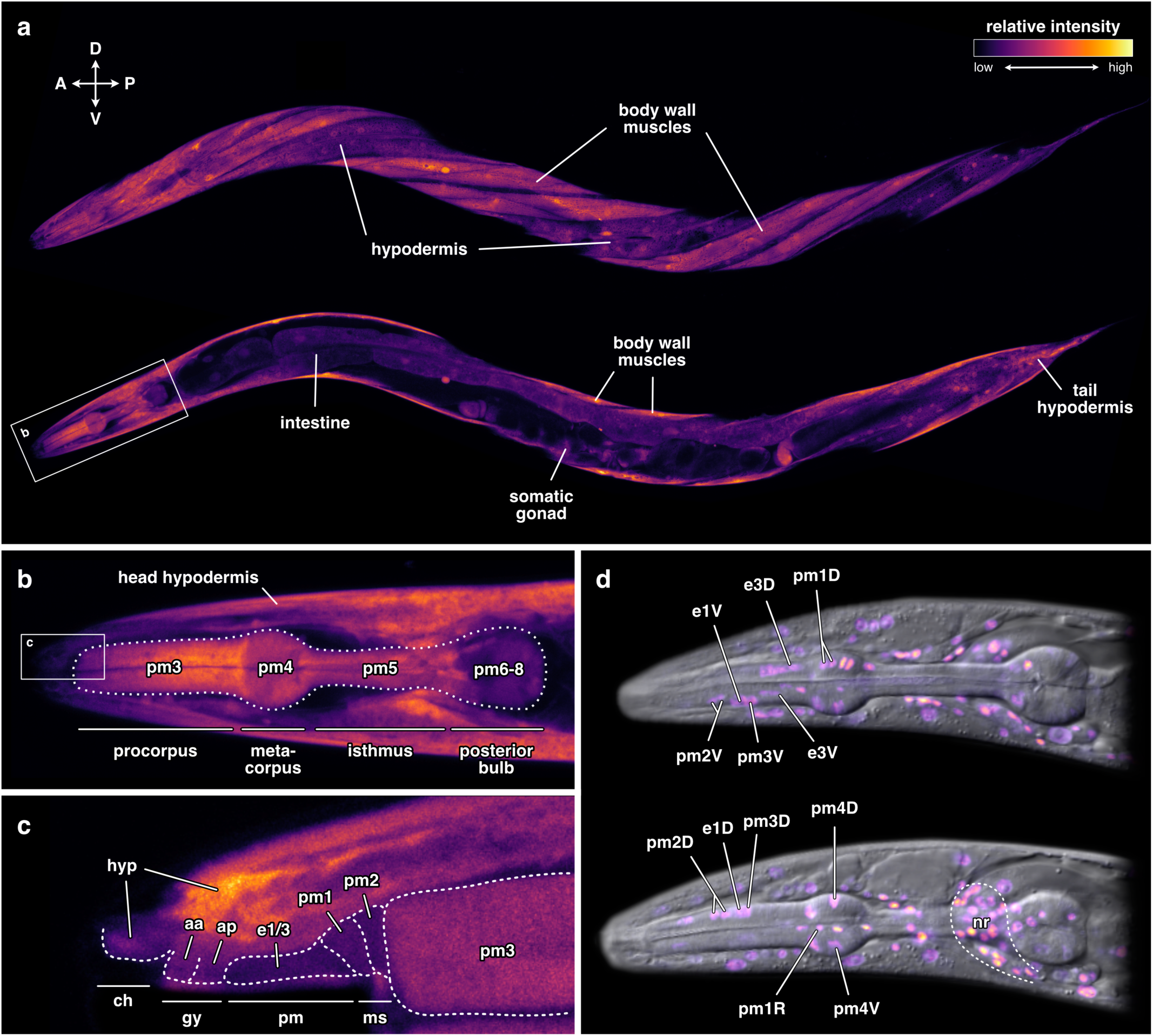
Expression and protein localization patterns of NHR-1 in *C. elegans*. (a) Pattern of *Cel-nhr-1* transcription in a young adult hermaphrodite. The same worm is depicted twice: top image shows its left lateral body surface; bottom image shows its mid-sagittal plane (A = anterior, P = posterior, D = dorsal, V = ventral). A zoom-in on the head region of the worm is depicted in (a) is provided in (b), showing the expression of *Cel-nhr-1* in the major pharyngeal muscle cells, as well as the head hypodermis. A close-up of the dorsal half of the stomatal tissues in another young hermaphrodite of the transcriptional reporter strain (c) reveals *Cel-nhr-1* expression in all of the cells which secrete the cuticular feeding structures (hyp = hypodermis, aa = anterior arcade syncytium, ap = posterior arcade syncytium, e1/3 = pharyngeal epithelial cells 1 and 3, pm = pharyngeal muscle cell, ch = cheilostom, gy = gymnostom, pm = promesostegostom, ms = metastegostom). A translational reporter strain (d) reveals that the *Cel-*NHR-1 protein localizes to the nuclei of the stomatal cells (D = dorsal, V = ventral, nr = nerve ring). Strength of fluorescence is indicated via a relative-intensity scale (a).

Taken together, the expression and protein-localization patterns of *Cel-*NHR-1 are highly consistent with the ones previously reported for *Cel-*NHR-40. Most importantly, both receptors are co-expressed in the anterior pharyngeal muscle cells in both *P. pacificus* and *C. elegans*, suggesting that this expression pattern was indeed already established in their last common ancestor. This, together with the fact that NHR-1 and NHR-40 co-regulate mouth development in *P. pacificus*, suggests that these receptors may in fact govern similar processes in *C. elegans*. To investigate this possibility, we created knock-out alleles for both of these receptors to see if that causes altered mouth morphologies reminiscent of those induced by the same kind of mutations in *P. pacificus* (Kieninger *et al*., 2016; Sieriebriennikov *et al*., 2020).

### Loss of *nhr-1* or *nhr-40* does not disrupt mouth morphogenesis in *C. elegans*

We obtained multiple independent homozygous mutant alleles of *Cel-nhr-1* and *Cel-nhr-40* by using CRISPR/CAS9 to introduce frameshift mutations into their gene bodies, which truncate the receptor proteins at the site of their DNA-binding domain (DBD) and render them non-functional. All of these homozygous mutants were viable, fertile, and showed no obvious morphological alterations. However, as the mouths of these nematodes are only a few micrometers long and wide (Figure 1a), which makes the identification of potential structural alterations difficult, we used landmark-based geometric morphometrics to quantify potential mutant phenotypes in mouth morphology (Figure 3c) (Theska *et al*., 2020). Two frameshift alleles, *Cel-nhr-1(tu1424)* and *Cel-nhr-40(tu1464)*, were chosen as reference alleles and used for all subsequent experiments and analyses (Figure 3a,b).

**Figure 3.**
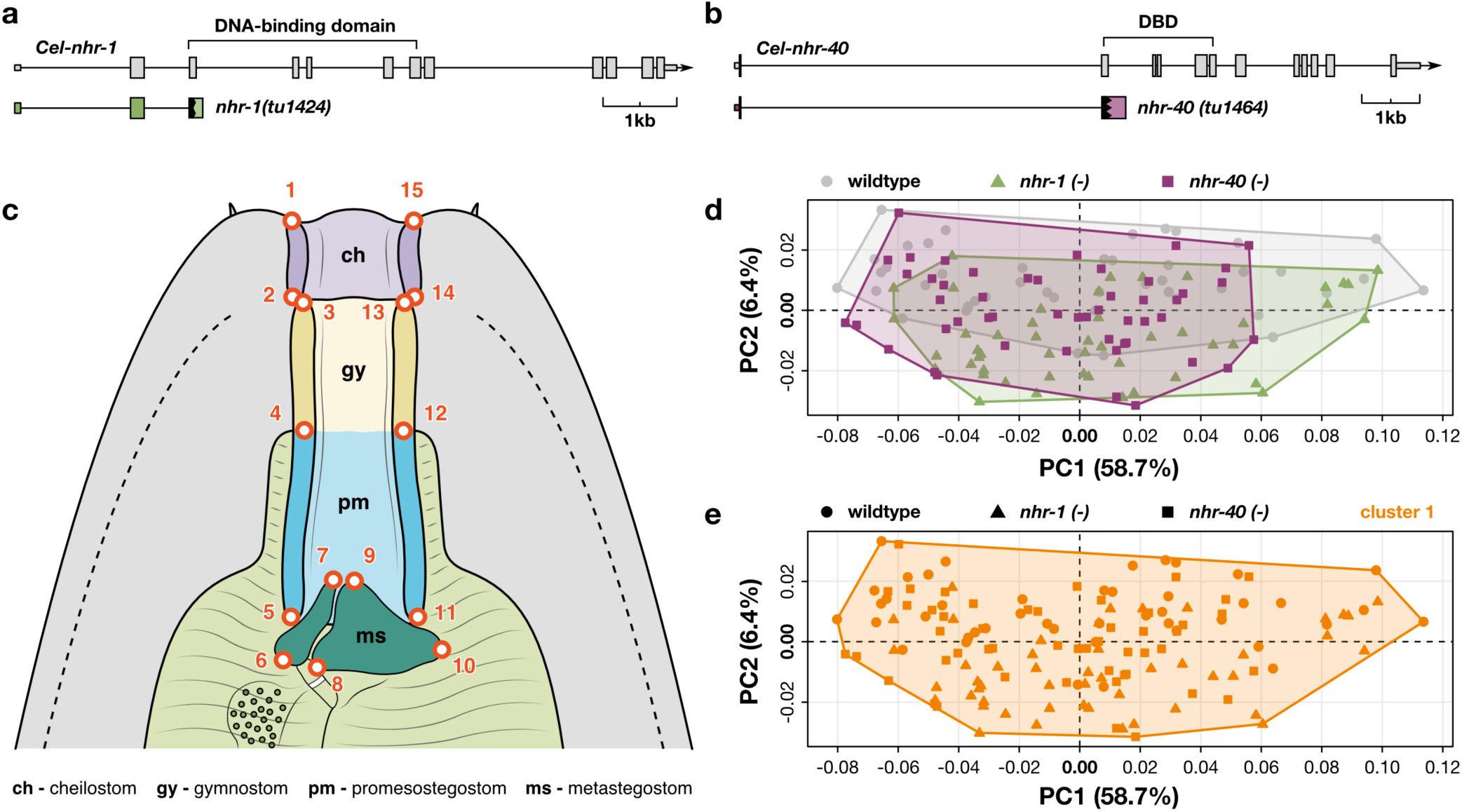
Neither the loss of NHR-1 or NHR-40 causes malformations in the *C. elegans* buccal capsule. (a, b) Gene models of *Cel-nhr-1* and *Cel-nhr-40* and their reference frameshift alleles which led to an early truncation of their DNA-binding domains (DBD). (c) Schematic representation of the adult *C. elegans* mouth depicting the landmark configuration used for geometric morphometric and clustering analysis (d, e). (d) PCA of form differences in wild-type *C. elegans*, as well as *Cel-nhr-1(tu1424)* and *Cel-nhr-40(1464)* mutants. Note that the ranges of morphological variation (outlined by convex hulls) in mutant strains strongly overlap with the wild-type range of morphological variation. (e) Unsupervised (model-based) clustering reveals that all 143 individuals belong to a single morphological cluster, demonstrating that mutant worms cannot be differentiated from wild-type worms based on their mouth morphology.

Surprisingly, we found that the loss of neither of these two NHRs affects the overall morphology of the adult mouth. A principal component analysis (PCA) of mouth-form data from 143 individuals revealed that, in the morphospace produced by PC1 and PC2 (which capture the majority of form variation in the data), wild-type worms and *nhr* mutants show homogenous and strongly overlapping ranges of morphological variation in their mouths (Figure 3d). This finding corroborated the superficial observation that wild-type and mutant worms were indistinguishable based on their morphology. We were able to estimate that PC1 is the only “meaningful” principal component (*sensu* Schlager, 2017) and that the other PCs are likely to contain noise rather than signal. Subsequent model-based clustering of all individuals based on the morphological variation contained in PC1 revealed that neither *Cel-nhr-1(tu1424)* nor *Cel-nhr-40(tu1464)* mutants could be classified as such or differentiated from wild-type worms in terms of their mouth morphology (Figure 3e) (note that this also holds true if clustering is perform based on the variation in PC2, or PC1 and PC2 together). Thus, geometric morphometric and model-based clustering analyses coherently suggest the absence of discernible malformations in the mouths of the two *nhr-*mutant strains when compared to wild-type animals (Figure 3d,e).

These findings are in contrast with the phenotypes caused by the loss of the same receptors in *P. pacificus,* which either lead to blatant structural malformations in the adult mouth or affect which of the worm’s alternative mouth morphologies is developmentally executed (Sieriebriennikov *et al*., 2020). Thus, the results of our morphological analyses in *C. elegans* did not support the hypothesis that NHR-1 and NHR-40 have a conserved biological function in the regulation of mouth morphogenesis across diplogastrids and “rhabditids”. Given that both receptors are deeply conserved between these worms and actively expressed in the same tissues, we wondered whether the absence of mouth-form related phenotypes in *Cel-nhr-1(tu1424)* and *Cel-nhr-40(tu1464)* mutants might be explained by differences in the regulatory targets of these receptors between the two species.

### NHR-1 and NHR-40 have largely different regulatory targets between *P. pacificus* and *C. elegans*

First, we wondered whether NHR-1 and NHR-40 have any common regulatory targets in *C. elegans*, just as they do in *P. pacificus*. To determine this, we first reanalyzed the transcriptomes of the *Ppa-nhr-1(tu1163)*, *Ppa-nhr-1(tu1164), Ppa-nhr-40(tu1418)*, and *Ppa-nhr-40(tu1423)* mutants obtained by Sieriebriennikov *et al*. (2020) alongside the transcriptomes of our *Cel-nhr-1(tu1424)* and *Cel-nhr-40(tu1464)* mutants. The purpose of the reanalysis of the *P. pacificus* data was two-fold: we wanted to analyze *nhr-*mutant transcriptomes in both species with the exact same pipeline and apply more stringent criteria in our differential expression analyses than in previous studies (e.g. including minimum effect sizes) (Figure 4 and Supplementary Figure 1).

**Figure 4.**
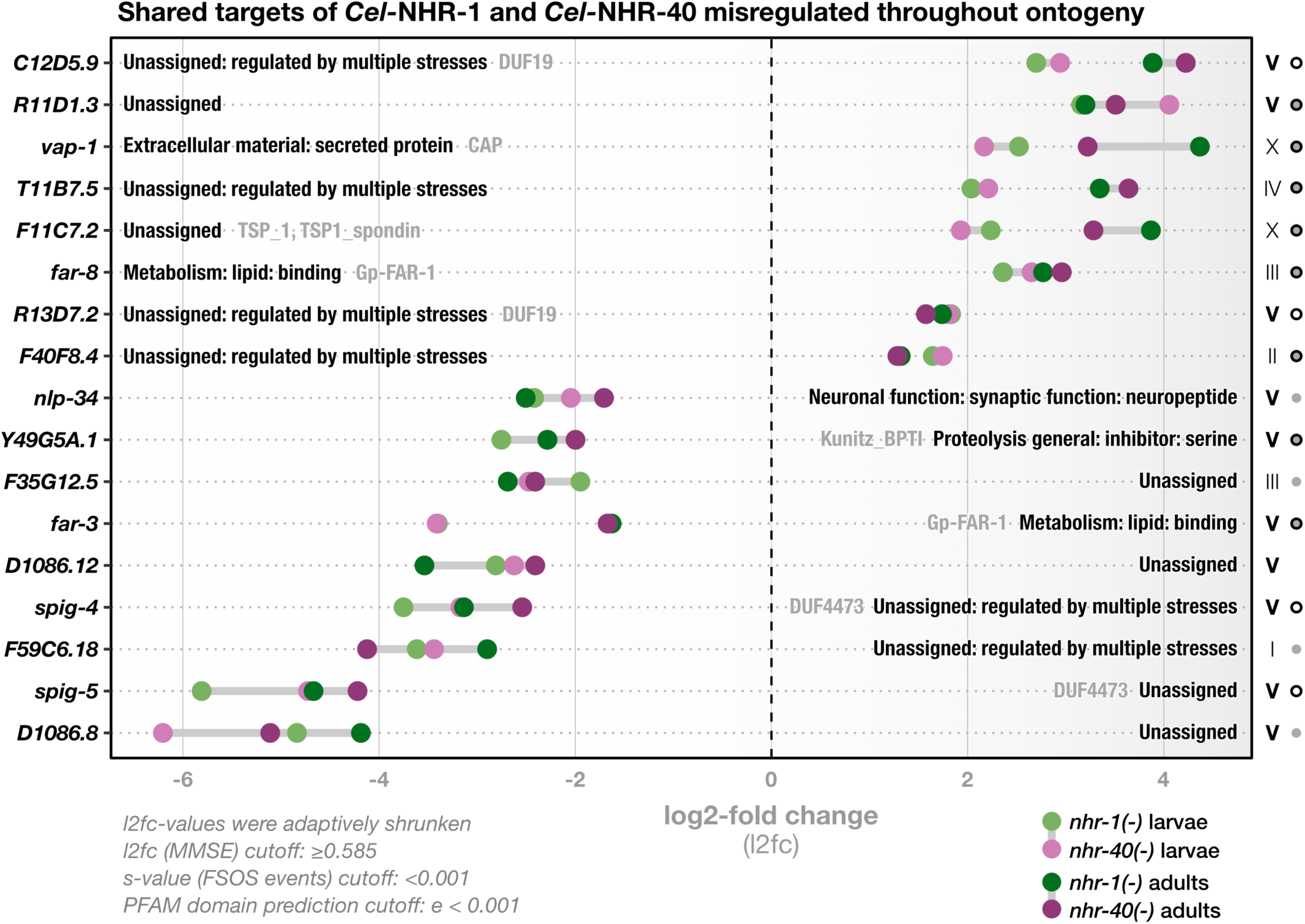
NHR-1 and NHR-40 share a pool of regulatory targets with currently unassigned biological functions. Bold names along the y-axis correspond to names of the shared regulatory targets of *Cel-*NHR-1 and *Cel-*NHR-40 which are misregulated throughout all developmental stages. Labels within the plot indicate functional information for the gene in form of WormCat terms (black) and/or predicted PFAM domains (gray). Roman numerals to the right of the plot indicate the chromosomal location of the shared targets. Note that the majority of target genes is located on chromosome V (bold letters indicate statistically overrepresented chromosomal locations). Dots to the right of the roman numerals indicate tissues in which the genes are known to be expressed (gray dot = neurons, black circle = glia cells, gray dot with black outline = neurons and glia cells). Cutoffs used in the differential gene expression analysis with DESeq2 and for PFAM domain prediction with HMMER are indicated in the lower left. MMSE = minimum mean squared error, FSOS = false sign or smaller.

The reanalysis of the *P. pacificus* mutant data largely corroborated the original findings and removed only a few putative targets. Sieriebriennikov *et al*. (2020) found, amongst a few others, 12 metallopeptidases (astacins), five secreted proteins (CAP-domain proteins), and two chitinases to be the major common regulatory targets of *Ppa-*NHR-1 and *Ppa-*NHR-40. Our reanalysis also recovered 12 metallopeptidases (Supplementary Figure 1) of which 10 were among the previously reported; all of them were found to be down-regulated in larval and adult worms. We also recovered the two chitinases and one of the secreted proteins among the common targets that are consistently mis-regulated throughout development (Supplementary Fig 1). Thus, this reanalysis of the *P. pacificus* transcriptome data both validated our computational pipeline and demonstrated that the original results by Sieriebriennikov *et al*. (2020) were robust against more stringent analytical cutoffs.

By performing similar analyses on the *C. elegans* transcriptome data we found that, just like in *P. pacificus*, *Cel-*NHR-1 and *Cel-*NHR-40 do indeed share a group of regulatory targets (Figure 4). A total of 17 genes are constitutively mis-regulated in both larvae and adults of *Cel-nhr-1(tu1424)* and *Cel-nhr-40(tu1464)* mutants, of which nine were down-regulated and eight up-regulated compared to wild-type worms. Strikingly, we found that the compositional patterns of the shared regulatory targets of these receptors is highly species-specific (Figure 4 and Supplementary Figure 1).

First, the comparative analysis revealed that targets which are shared by NHR-1 and NHR-40 in a stage-independent manner were biased towards a specific chromosome in either species. Where they were biased towards the X chromosome in *P. pacificus* (*P* = 1.5e-07, Supplementary Fig 1), they were biased towards chromosome V in *C. elegans* (*P* = 0.00011, Figure 4). These chromosomes are non-homologous (Yoshida *et al*., 2023).

Second, we did not find a single metallopeptidase or chitinase among the stage-independently shared targets of NHR-1 and NHR-40 in *C. elegans*. Thus, with the exception of a single CAP-domain-containing secreted protein (*vap-1*), none of the protein classes which were found to constitute the major regulatory targets of both receptors in *P. pacificus* could be found in a similar pool of targets for same receptors in *C. elegans*. Intriguingly, we found that 12 of the 17 shared regulatory targets of NHR-1 and NHR-40 in the latter species currently have no assigned biological roles, although six of them are at least known to be regulated by various stress factors (Figure 4). We wondered whether any of these biologically uncharacterized genes encode known protein family (pfam) domains (Finn *et al*., 2016), which could at least hint at their biochemical functions. We were able to predict domains in the amino acid sequences encoded by only five of the biologically uncharacterized genes, and just a single one of them - a thrombospondin (*F11C7.2*) - contains a domain that is somewhat understood. In contrast, the other four domain-containing proteins possess seemingly nematode-specific domains of unknown function (DUF19 and DUF4473). Therefore, most of NHR-1’s and NHR-40’s shared regulatory targets in *C. elegans* currently remain uncharacterized both at the biological and biochemical levels. Amongst the few shared targets for which a biological function is already known, we found two lipid-binding proteins with metabolic roles (*far-3* and *far-8*), a neuropeptide-like protein (*nlp-34*), and a serine protease inhibitor (*Y49G5A.1*) (Figure 4). Seemingly, these genes are not easily grouped into any overarching biological process that might be co-regulated by NHR-1 and NHR-40 in this species. Yet, all but one of their shared regulatory targets are known to be expressed predominantly in glia cells, sensory neurons, interneurons, motoneurons, or combinations thereof (Figure 4) (Taylor *et al*., 2021). Therefore, we speculate that, by co-regulating the neuronal expression of a pool of shared targets with hitherto unassigned biological roles, NHR-1 and NHR-40 might jointly control the execution of a neuronal (and possibly sensory) response in *C. elegans*.

Third, stage-specific overrepresentation analyses for functional categories amongst the identified target genes of NHR-1 and NHR-40 revealed that these receptors control essentially the same kind of regulatory targets (metallopeptidases, chitinases, and secreted proteins) throughout the development of *P. pacificus* and that only a few targets are specific to either a single receptor or developmental stage (Figure 5). Interestingly, the same does not hold true for *C. elegans*. In fact, the genes with currently unassigned biological functions were the only category that was overrepresented among the targets of both mutants in a stage-independent manner (Figure 5). These genes aside, we found multiple stage-specific common targets of NHR-1 and NHR-40 in *C. elegans*. For example, approximately half of all genes which encode major sperm proteins are down-regulated in larvae, and more than a fifth of all collagen-encoding genes were down-regulated in adults of both *nhr* mutants. Other adult-specific common targets include hedgehog-like proteins, acid phosphatases involved in lysosomal protein breakdown, and functionally uncharacterized prions. Almost all of these targets were consistently down-regulated in both mutant strains. The only adult-specific group of shared putative targets that is consistently up-regulated are CUB-domain proteins (Figure 5), which play important roles in the worm’s innate immune responses to pathogenic bacteria and fungi (Holdorf *et al*., 2020). Furthermore, our analyses revealed multiple receptor-specific target groups. In adult *Cel-nhr-1(tu1424)* mutants, broad aspects of amino acid and lipid metabolism are mis-regulated. This includes amino-acid breakdown, lipolysis, fatty-acid synthesis, and β-oxidation. Similarly affected are genes encoding enzymes with broad metabolic roles like flavin-containing monooxygenases and short chain dehydrogenases (Figure 5). Additionally, various stress reactions such as detoxification via p450 cytochromes and UDP-glucuronossyltransferases, immune responses involving C-type lectins, and unfolded protein responses in the endoplasmic reticulum seem to require NHR-1, too (Figure 5). NHR-40, on the other hand, seems to regulate some proteolytic processes in adult worms, including (surprisingly) a small number of metallopeptidases - the main targets of both NHR-1 and NHR-40 in *P. pacificus* (Figure 5).

**Figure 5.**
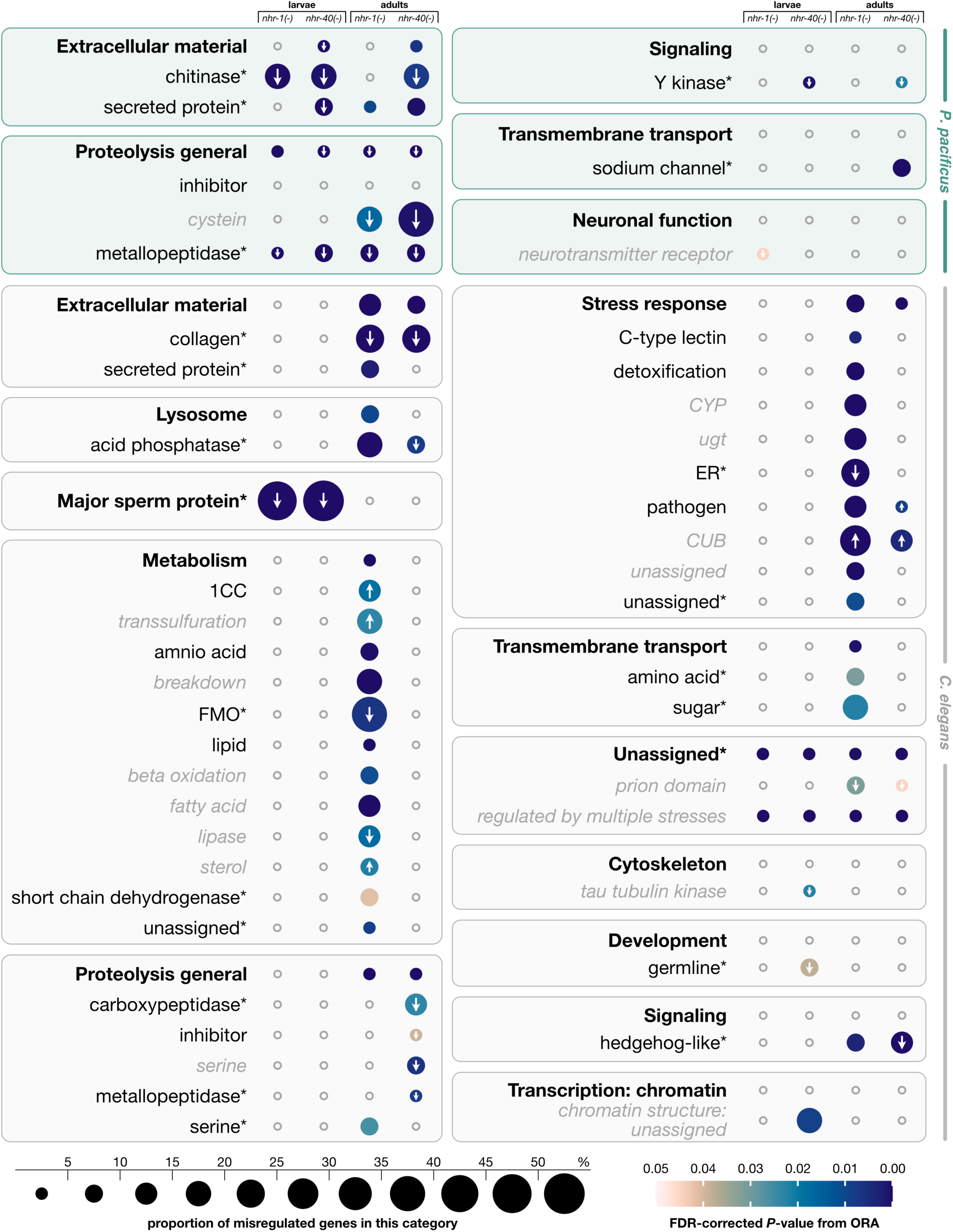
Comparative transcriptomics of *nhr-1* and *nhr-40* mutants in *P. pacificus* and *C. elegans* reveals species-specific composition of regulatory targets. Bubble chart showing overrepresented functional categories among the targets of each receptor in larvae and/or adults of *C. elegans* and *P. pacificus*. Enriched categories were broken down hierarchically. Boxes represent functional categories for which any overrepresentation signal could be detected. Functional categories were broken down into three hierarchical levels of specificity. Bold black font indicates the highest functional level (i.e., the least specific) and the name of the functional category represented by the box. Non-bold black font indicates the second category level (i.e., a more specific aspect of the overarching category). Gray italicized font indicates the most specific (and least inclusive) functional terms within the overarching category, for which overrepresentation signals could be detected. Asterisks (*) next to a category term indicate that no more-specific lower-level categories are defined within this term. The presence of bubbles indicates overrepresentation of the given functional category among target genes; absence of bubbles (depicted as gray circles) indicates no overrepresentation. Color bar indicates the statistical support for the overrepresentation signal (FDR-corrected *P-*values were derived from Fisher’s Exact test). Bubble size indicates the percentage of genes in the given functional category which are misregulated in that particular sample type (out of all genes in the genome with that particular functional annotation). White arrows indicate that all of these targets are consistently up- or down-regulated. FDR = false discovery rate, ORA = overrepresentation analysis.

## Discussion

In this study, we took a comparative evo-devo approach to elucidate the evolutionary history of a GRN that underlies the development of novel feeding structures in diplogastrid nematodes. Using *C. elegans* as a representative member of “rhabditids” (the paraphyletic outgroup of flap-carrying nematodes from which the Diplogastridae emerged), we investigated whether the mouth-morphogenesis related functions of NHR-1 and NHR-40 (Figure 1f) predated the emergence of diplogastrid teeth, or whether they evolved in conjunction with them (Figure 1e).

We anticipated to find that these two transcription factors have a conserved function in feeding-structure development between *C. elegans* and *P. pacificus*, based on a combination of observations: First, the teeth of diplogastrids are individualized novelties that evolved from “rhabditid” flaps. As such, both of these feeding structures can be homologized and traced back to the metastegostom of their last common ancestor (LCA). Second, the two molecular players of the phenotype-execution module, which are involved in cuticle synthesis, DPY-6 and CHS-1, have already been found to exert conserved functions in *C. elegans* and *P. pacificus* (Sun & Theska *et al*., 2022; Sun *et al*., 2023), further corroborating the hypothesis that this module was established before the flap-to-tooth transformation. Lastly, the two NHRs, which control the aspect of the phenotype-execution module that governs cuticle modification and degradation, are deeply conserved between *C. elegans* and *P. pacificus*, too (Sieriebriennikov *et al*., 2020), and thus likely to regulate similar developmental processes in both species.

However, the results of the integrative analysis described in this study do not support our initial expectations. Despite the fact that the expression studies on *Cel-* NHR-1, together with the available expression data on *Cel-*NHR-40, indicate that both receptors are co-expressed in the tissues which give rise to the feeding structures of both *C. elegans* and *P. pacificus*, genetic inactivation of either receptor does not disrupt or otherwise affect feeding-structure development in the “rhabditid” species. Therefore, feeding-structure development in *C. elegans* and *P. pacificus* is not controlled by the exact same genetic module. This observation was further supported by the comparative transcriptomic experiments, which revealed that NHR-1 and NHR-40 have highly species-specific types of regulatory targets and thus control different biological processes in *C. elegans* and *P. pacificus*.

Throughout *P. pacificus’s* ontogeny, both receptors co-regulate the expression of a narrow range of “core targets”, which is essentially restricted to the chitinases, metallopeptidases, and secreted proteins involved in the cuticle-modifying and -degrading processes of feeding-structure development. The same receptors also share numerous regulatory targets in *C. elegans*, but, as opposed to their *P. pacificus* counterparts, they represent functionally more diverse groups of genes that are often co-regulated in a stage-specific manner (Figure 5). Additionally, in *C. elegans*, both receptors regulate various non-overlapping biological processes. Interestingly, none of *C. elegans’s* chitinase genes were affected by the loss of either receptor at any stage of development. Yet, we found that five secreted proteins displayed altered expression levels in adult *Cel-nhr-1(tu1424)* mutants (of which one is an ortholog to a core target of *Ppa-*NHR-1 and *Ppa-*NHR-40), and that five metallopeptidases were misregulated in adult *Cel-nhr-40(tu1464)* mutants (of which one is an ortholog to two of *Ppa-*NHR-1’s and *Ppa-*NHR-40’s core targets). Besides the differences between the NHR-1 and NHR-40 targets in these two species, there are some correlations of currently unknown significance. For example, independent of the developmental stage, both receptors co-regulate the expression of small pools of genes that tend to be located on a particular chromosome (X in *P. pacificus* vs. V in *C. elegans*) and co-expressed in specific cell types (pharyngeal gland in *P. pacificus* vs. glia and neurons in *C. elegans*). Furthermore, and adding to previous reports (Sieriebriennikov *et al*., 2020), we found that the expression levels of either receptor remained unaffected by the inactivation of the other, at any stage of development in both species. This suggests that, within their respective GRNs, NHR-1 and NHR-40 do not activate or repress each other transcriptionally, but that they might co-regulate species-specific pools of shared target genes via post-transcriptional interactions, possibly by binding to the same promotor regions as heterodimers.

Taken together, the findings presented in this study could support two different evolutionary scenarios. Either, the phenotype-execution module which governs feeding-structure morphogenesis in diplogastrids indeed already existed in the LCA of *C. elegans* and *P. pacificus.* In this scenario, NHR-1 and NHR-40 used to control the expression of many metallopeptidases, chitinases, and secreted proteins in order to modify feeding structures throughout the ontogeny of “rhabditids” and diplogastrids. This, in turn, would mean that the entire phenotype-execution module (in its diplogastrid-like form) already existed before the Diplogastridae and their teeth evolved, and that *C. elegans* happens to be a “rhabditid” species in which this module secondarily disintegrated as NHR-1 and NHR-40 acquired new regulatory targets. In this case, the small number of metallopeptidases and secreted proteins we found to be misregulated in adults of either *Cel-nhr-1(tu1424)* or *Cel-nhr-40(tu1464)* mutants may reflect minor remnants of this ancestral genetic module. Alternatively, the phenotype-execution module of *P. pacificus’s* feeding-structure GRN assembled and gained its function in conjunction with the emergence of Diplogastridae. In this scenario, the absence of discernible mouth-morphogenesis related functions for the two NHRs in *C. elegans* would be explained by the fact these receptors used to control different biological processes in the LCA of “rhabditids” and diplogastrids (probably ones akin to the metabolic and physiological processes we found to be controlled by them in *C. elegans*). However, final conclusions regarding the ancestral regulatory functions of NHR-1 and NHR-40 cannot yet be drawn, since our study only addressed their roles in *C. elegans.* In order to established character polarity for the regulatory functions of these receptors (as they differ so clearly between the two species for which they are now known), future studies will have to determine the targets of NHR-1 and NHR-40 in additional “rhabditid” species more closely and more distantly related to the Diplogastridae than *C. elegans*. This way, we will learn where the phenotype-execution module of the diplogastrids emerged, and whether *Caenorhabditis* nematodes secondarily lost this module or never possessed it in the first place. This, however, is complicated by the fact that it is currently unclear which of the “rhabditids” constitutes the sister taxon of Diplogastridae, even though current studies suggest that it is *Rhabditoides* (van Megen *et al*., 2009; Sudhaus, 2014).

In summary, the currently available data show that the phenotype-execution module that underlies feeding-structure development in diplogastrids is not functionally conserved between *P. pacificus* and *C. elegans*, suggesting that it was subject to developmental systems drift (Sommer, 2012; Haag *et al*., 2018) during the flap-to-tooth transformation that accompanied the “rhabditid”-to-diplogastrid transition. We speculate that the two ancient transcription factors NHR-1 and NHR-40 were likely co-opted into a new regulatory context when the diplogastrid teeth evolved, given that most of the genes in the feeding-structure GRN (including many of NHRs’ shared targets) derived from diplogastrid-specific gene duplication events (Sieriebriennikov *et al.,* 2020). In any case, the numerous details surrounding the evolutionary history of the flap-to-tooth transformation in diplogastrids highlight the complexity of the developmental genetic processes which underlie morphological evolution (Carroll, 2008), and are in line with similarly complex evolutionary histories identified in other evo-devo systems, such as the evolution of novel extraembryonic tissues in insects (van der Zee *et al*., 2005; Panfilio *et al*., 2013).

## Acknowledgements

The plasmid pDD268 was a gift from Daniel Dickinson and Bob Goldstein (Addgene plasmid #132523; http://n2t.net/addgene:132523; RRID:Addgene_132523). We thank the Light Microscopy facility at the Max Planck Institute for Biology, for their assistance with cLSM. We are grateful to Heike Haussmann for freezing samples of all the worm strains produced for this study. Ziduan Han and Bogdan Sieriebriennikov provided expertise on molecular biological approaches in early stages of this project. Additionally, we thank Christian Rödelsperger, Devansh R. Sharma, Michael S. Werner, and James Lightfoot for helpful discussions of our work. We are grateful to Tess Renahan for critically reading this manuscript and for valuable suggestions throughout all stages of the project. T.T. was supported by the International Max Planck Research School (IMPRS) ’From Molecules to Organisms’.

## Data availability

Raw reads from the mRNA-seq experiments have been deposited in the European Nucleotide Archive (ENA) under the accession number PRJEB68220. Landmark data for geometric morphometrics, results of the differential expression analyses, and results of the overrepresentation analyses are provided in the supplementary material (Supplementary Data 1-5).

---

**Supplementary Figure 1.**
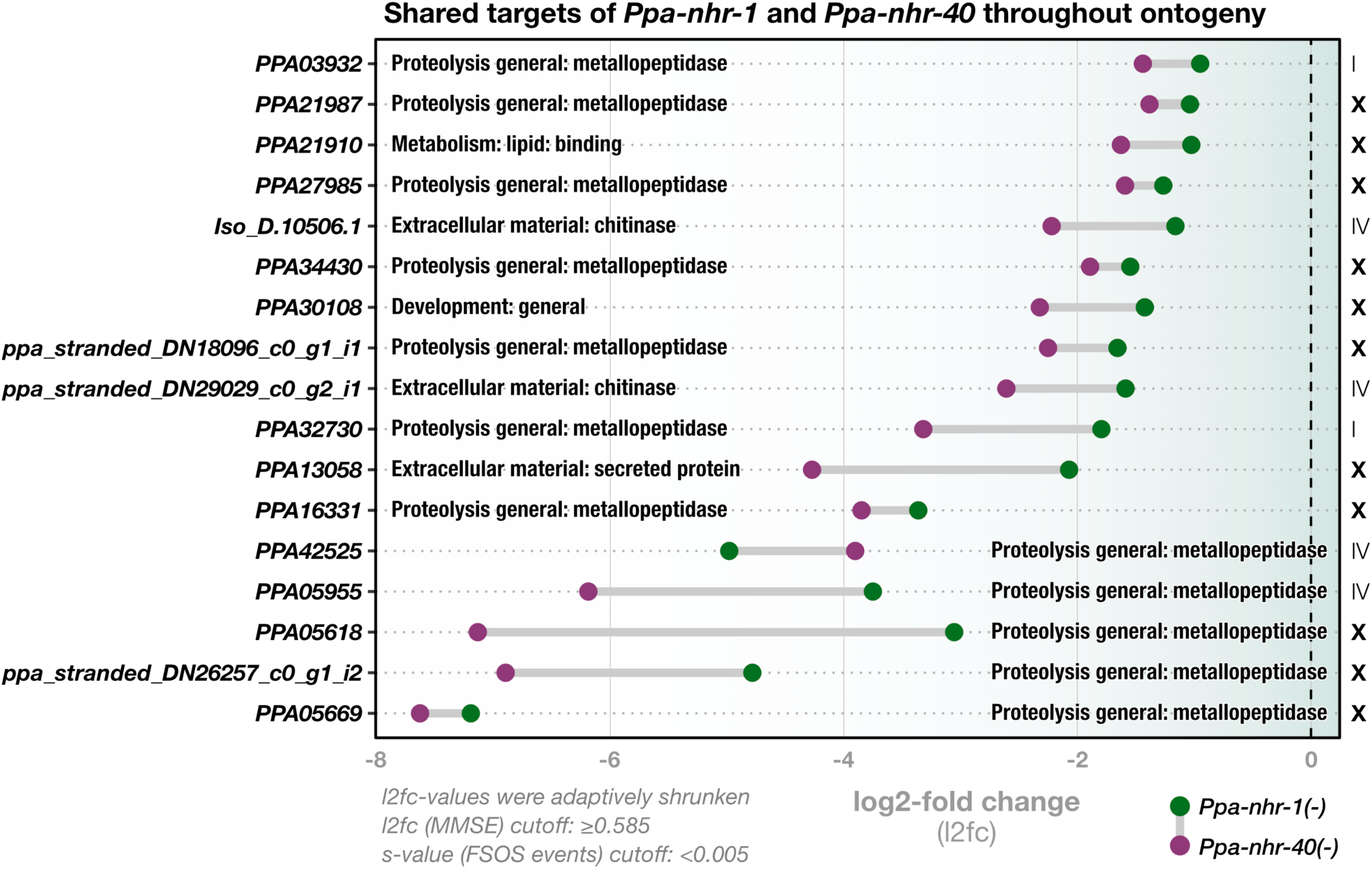
Re-analysis of target genes that are constitutively misregulated in *Ppa-nhr-1* and *Ppa-nhr-40* mutants. Bold names along the y-axis correspond to names of the shared regulatory target genes of *Ppa-*NHR-1 and *Ppa-* NHR-40 which are misregulated throughout ontogeny. Labels within the plot indicate functional information for the gene in form of WormCat terms. Roman numerals to the right of the plot indicate the chromosomal location of the shared targets. Note that the majority of the targets is located on the X chromosome (bold letters indicate statistically overrepresented chromosomal location). Cutoffs used in the differential gene expression analysis with DESeq2 are indicated in the lower left. MMSE = minimum mean squared error, FSOS = false sign or smaller.

## References

1. Adams, D. C., & Otárola-Castillo, E. (2013). geomorph: An R package for the collection and analysis of geometric morphometric shape data. Methods in Ecology and Evolution, 4(4), 393–399.

2. Baken, E. K., Collyer, M. L., Kaliontzopoulou, A., & Adams, D. C. (2021). Geomorph v4. 0 and gmShiny: Enhanced analytics and a new graphical interface for a comprehensive morphometric experience. Methods in Ecology and Evolution, 12(12), 2355–2363.

3. Brožová, E., Šimečková, K., Kostrouch, Z., Rall, J. E., & Kostrouchová, M. (2006). NHR-40, a *Caenorhabditis elegans* supplementary nuclear receptor, regulates embryonic and early larval development. Mechanisms of Development, 123(9), 689–701.

4. Burr, A., & Baldwin, J. G. (2016). The nematode stoma: Homology of cell architecture with improved understanding by confocal microscopy of labeled cell boundaries. Journal of Morphology, 277(9), 1168–1186.

5. Carroll, S.B. (2008). Evo-devo and an expanding evolutionary synthesis: a genetic theory of morphological evolution. Cell, 134, 25–36.

6. Corsi, A. K., Wightman, B., & Chalfie, M. (2015). A transparent window into biology: A primer on *Caenorhabditis elegans*. Genetics, 200(2), 387–407.

7. De Ley, P., Van de Velde, M.C., Mounport, D., Baujard, P. & Cooman, A. (1995) Ultrastructure of the stoma in Cephalobidae, Panagrolaimidae and Rhabditidae, with a proposal for a revised stoma terminology in Rhabditida (Nematoda). Nematologica, 41(1-4), 153–182.

8. Dickinson, D. J., Pani, A. M., Heppert, J. K., Higgins, C. D., & Goldstein, B. (2015). Streamlined genome engineering with a self-excising drug selection cassette. Genetics, 200(4), 1035–1049.

9. Emms, D. M., & Kelly, S. (2019). OrthoFinder: Phylogenetic orthology inference for comparative genomics. Genome Biology, 20, 1–14.

10. Finn, R. D., Coggill, P., Eberhardt, R. Y., Eddy, S. R., Mistry, J., Mitchell, A. L., Potter, S. C., Punta, M., Qureshi, M., & Sangrador-Vegas, A. (2016). The Pfam protein families database: Towards a more sustainable future. Nucleic Acids Research, 44(D1), D279–D285.

11. Fürst von Lieven, A., & Sudhaus, W. (2000). Comparative and functional morphology of the buccal cavity of Diplogastrina (Nematoda) and a first outline of the phylogeny of this taxon. Journal of Zoological Systematics and Evolutionary Research, 38(1), 37–63.

12. Garnier, S., Ross, N., Rudis, R., Camargo, P. A., Sciaini, M., & Scherer, C. (2021). viridis—Colorblind-friendly color maps for R. R Package Version 0.6, 1.

13. Gentleman, R. C., Carey, V. J., Bates, D. M., Bolstad, B., Dettling, M., Dudoit, S., Ellis, B., Gautier, L., Ge, Y., & Gentry, J. (2004). Bioconductor: Open software development for computational biology and bioinformatics. Genome Biology, 5(10), 1–16.

14. Ghanta, K. S., Ishidate, T., & Mello, C. C. (2021). Microinjection for precision genome editing in Caenorhabditis elegans. STAR Protocols, 2(3), 100748.

15. Haag, E. S., Fitch, D. H., & Delattre, M. (2018). From “the worm” to “the worms” and back again: The evolutionary developmental biology of nematodes. Genetics, 210(2), 397–433.

16. Harry, C. J., Messar, S. M., & Ragsdale, E. J. (2022). Comparative reconstruction of the predatory feeding structures of the polyphenic nematode Pristionchus pacificus. Evolution & Development, 24(1–2), 16–36.

17. Holdorf, A. D., Higgins, D. P., Hart, A. C., Boag, P. R., Pazour, G. J., Walhout, A. J., & Walker, A. K. (2020). WormCat: An online tool for annotation and visualization of *Caenorhabditis elegans* genome-scale data. Genetics, 214(2), 279– 294.

18. Howard, R.J., Giacomelli, M., Lozano-Fernandez, J., Edgecombe, G.D., Fleming, J.F., Kristensen, R.M., … & Pisani, D. (2022). The Ediacaran origin of Ecdysozoa: integrating fossil and phylogenomic data. Journal of the Geological Society, 179(4):jgs2021–2107.

19. Kieninger, M. R., Ivers, N. A., Rödelsperger, C., Markov, G. V., Sommer, R. J., & Ragsdale, E. J. (2016). The nuclear hormone receptor NHR-40 acts downstream of the sulfatase EUD-1 as part of a developmental plasticity switch in *Pristionchus*. Current Biology, 26(16), 2174–2179.

20. Love, M. I., Anders, S., Kim, V., & Huber, W. (2015). RNA-Seq workflow: Gene-level exploratory analysis and differential expression. F1000Research, 4, 1070.

21. Love, M. I., Huber, W., & Anders, S. (2014). Moderated estimation of fold change and dispersion for RNA-seq data with DESeq2. Genome Biology, 15(12), 1–21.

22. Malakhov, V.V. (1994). Nematodes: Structure, development, classification and phylogeny. Smithsonian Books.

23. Minelli, A. (2015). Grand challenges in evolutionary developmental biology. Frontiers in Ecology and Evolution, 2, 85, 1–11.

24. Mistry, J., Finn, R. D., Eddy, S. R., Bateman, A., & Punta, M. (2013). Challenges in homology search: HMMER3 and convergent evolution of coiled-coil regions. Nucleic Acids Research, 41(12), e121–e121.

25. Mitteroecker, P., Gunz, P., Bernhard, M., Schaefer, K., & Bookstein, F. L. (2004). Comparison of cranial ontogenetic trajectories among great apes and humans. Journal of Human Evolution, 46(6), 679–698.

26. Müller, G. B. (2021). Developmental Innovation and Phenotypic Novelty. Evolutionary Developmental Biology: A Reference Guide, Nuño de la Rosa, L., Müller, G.B. (Ed.) (pp.69–84). Springer.

27. Panfilio, K.A., Oberhofer, G., & Roth, S. (2013). High plasticity in epithelial morphogenesis durin insect dorsal closure. Biology Open, 2(11), 1108–1118.

28. Patro, R., Duggal, G., Love, M. I., Irizarry, R. A., & Kingsford, C. (2017). Salmon provides fast and bias-aware quantification of transcript expression. Nature Methods, 14(4), 417–419.

29. Pedersen, T. L., & Crameri, F. (2023). Scico: Colour palettes based on the scientific colour-maps. R Package Version 1.4.0.

30. R Core Team, R. (2023). *R: A language and environment for statistical computing*. R Foundation for Statistical Computing. https://www.R-project.org/

31. Riebesell, M., & Sommer, R.J. (2017). Three-dimensional reconstruction of the pharyngeal gland cells in the predatory nematode *Pristionchus pacificus*. Journal of Morphology, 278(12), 1656–1666.

32. Robinson-Rechavi, M., Maina, C. V., Gissendanner, C. R., Laudet, V., & Sluder, A. (2005). Explosive lineage-specific expansion of the orphan nuclear receptor HNF4 in nematodes. Journal of Molecular Evolution, 60, 577–586.

33. Schindelin, J., Arganda-Carreras, I., Frise, E., Kaynig, V., Longair, M., Pietzsch, T., Preibisch, S., Rueden, C., Saalfeld, S., & Schmid, B. (2012). Fiji: An open-source platform for biological-image analysis. Nature Methods, 9(7), 676–682.

34. Schlager, S. (2017). Morpho and Rvcg–Shape Analysis in R: R-Packages for geometric morphometrics, shape analysis and surface manipulations. In Zheng, G., Li, S., Szekely, G. (Ed.), Statistical Shape and Deformation Analysis (pp. 217–256). Elsevier.

35. Scrucca, L., Fop, M., Murphy, T. B., & Raftery, A. E. (2016). mclust 5: Clustering, classification and density estimation using Gaussian finite mixture models. The R Journal, 8(1), 289.

36. Sieriebriennikov, B., Prabh, N., Dardiry, M., Witte, H., Röseler, W., Kieninger, M. R., Rödelsperger, C., & Sommer, R. J. (2018). A developmental switch generating phenotypic plasticity is part of a conserved multi-gene locus. Cell Reports, 23(10), 2835–2843.

37. Sieriebriennikov, B., Sun, S., Lightfoot, J. W., Witte, H., Moreno, E., Rödelsperger, C., & Sommer, R. J. (2020). Conserved nuclear hormone receptors controlling a novel plastic trait target fast-evolving genes expressed in a single cell. PLoS Genetics, 16(4), e1008687.

38. Sommer, R. J. (2012). Evolution of regulatory networks: Nematode vulva induction as an example of developmental systems drift. In Soyer, O. (Ed.), Evolutionary Systems Biology (pp. 79–91). Springer.

39. Sommer, R. J. (Ed.) (2015). Pristionchus pacificus: A nematode model for comparative and evolutionary biology (Vol. 11). Leiden, The Netherland: Brill.

40. Sommer, R. J. (2020). Phenotypic plasticity: From theory and genetics to current and future challenges. Genetics, 215(1), 1–13.

41. Sommer, R. J., Dardiry, M., Lenuzzi, M., Namdeo, S., Renahan, T., Sieriebriennikov, B., & Werner, M. S. (2017). The genetics of phenotypic plasticity in nematode feeding structures. Open Biology, 7(3), 160332.

42. Srivastava, A., Malik, L., Sarkar, H., Zakeri, M., Almodaresi, F., Soneson, C., Love, M. I., Kingsford, C., & Patro, R. (2020). Alignment and mapping methodology influence transcript abundance estimation. Genome Biology, 21(1), 1–29.

43. Stephens, M. (2017). False discovery rates: A new deal. Biostatistics, 18(2), 275–294.

44. Stiernagle, T. (2006). Maintenance of *C. elegans*. In the *C. elega*ns research community (Ed.), WormBook: The Online Review of C. elegans Biology.

45. Sudhaus, W. (2014). 7.17 Order Rhabditina: “Rhabditidae”. In Schmidt-Rhaesa, A. (Ed.), Volume 2 Nematoda (pp. 537–556). Berlin, Boston: De Gruyter.

46. Sun, S., Theska, T., Witte, H., Ragsdale, E. J., & Sommer, R. J. (2022). The oscillating Mucin-type protein DPY-6 has a conserved role in nematode mouth and cuticle formation. Genetics, 220(3), iyab233.

47. Sun, S., Witte, H., & Sommer, R. J. (2023). Chitin contributes to the formation of a feeding structure in a predatory nematode. Current Biology, 33(1), 15–27.

48. Sural, S., & Hobert, O. (2021). Nematode nuclear receptors as integrators of sensory information. Current Biology, 31(19), 4361–4366.

49. Susoy, V., Ragsdale, E. J., Kanzaki, N., & Sommer, R. J. (2015). Rapid diversification associated with a macroevolutionary pulse of developmental plasticity. Elife, 4, e05463.

50. Taylor, S. R., Santpere, G., Weinreb, A., Barrett, A., Reilly, M. B., Xu, C., Varol, E., Oikonomou, P., Glenwinkel, L., & McWhirter, R. (2021). Molecular topography of an entire nervous system. Cell, 184(16), 4329–4347.

51. Theska, T., Sieriebriennikov, B., Wighard, S. S., Werner, M. S., & Sommer, R. J. (2020). Geometric morphometrics of microscopic animals as exemplified by model nematodes. Nature Protocols, 15(8), 2611–2644.

52. van der Zee, M., Berns, N., & Roth, S. (2005). Distinct functions of the *Tribolium zerknüllt* genes in serosa specification and dorsal closure. Current Biology, 15(7), 624–636.

53. van Megen, H., van den Elsen, S., Holterman, M., Karssen, G., Mooyman, P., Bongers, T., … & Helder, J. (2009). A phylogenetic tree of nematodes based on about 1200 full-length small subunit ribosomal DNA sequences. Nematology, 11(6), 927–950.

54. Wagner, G. P. (2014). Homology, Genes, and Evolutionary Innovation. Princeton University Press.

55. Werner, M. S., Claaßen, M. H., Renahan, T., Dardiry, M., & Sommer, R. J. (2018). Adult influence on juvenile phenotypes by stage-specific pheromone production. Iscience, 10, 123–134.

56. Werner, M. S., Loschko, T., King, T., Reich, S., Theska, T., Franz-Wachtel, M., Macek, B., & Sommer, R. J. (2023). Histone 4 lysine 5/12 acetylation enables developmental plasticity of *Pristionchus* mouth form. Nature Communications, 14(1), 2095.

57. Werner, M. S., Sieriebriennikov, B., Loschko, T., Namdeo, S., Lenuzzi, M., Dardiry, M., Renahan, T., Sharma, D. R., & Sommer, R. J. (2017). Environmental influence on *Pristionchus pacificus* mouth form through different culture methods. Scientific Reports, 7(1), 7207.

58. Wickham, H., Averick, M., Bryan, J., Chang, W., McGowan, L. D., François, R., Grolemund, G., Hayes, A., Henry, L., & Hester, J. (2019). Welcome to the Tidyverse. Journal of Open Source Software, 4(43), 1686.

59. Wright, K. (1976). Functional organization of the nematode’s head. The Organization of Nematodes, Neil A. Croll (Ed.). *Academic Press*, *London, UK*, 71–105.

60. Yoshida, K., Rödelsperger, C., Röseler, W., Riebesell, M., Sun, S., Kikuchi, T., & Sommer, R. J. (2023). Chromosome fusions repatterned recombination rate and facilitated reproductive isolation during *Pristionchus* nematode speciation. Nature Ecology & Evolution, 7(3), 424–439.

